# Identification of small regulatory RNAs involved in persister formation

**DOI:** 10.1101/310631

**Authors:** Shanshan Zhang, Shuang Liu, Nan Wu, Wenhong Zhang, Ying Zhang

## Abstract

Small regulatory RNA (srRNA) is widely distributed in three kingdoms of life and fulfills functions in many aspects of cellular life, but their role in bacterial persistence remains unknown. In this study, we comprehensively interrogated the expression levels of the known srRNAs on three critical time points, stage 1 (S1) where no persisters are formed, stage 2 (S2) where persisters are beginning to appear, and stage 3 (S3) where persister numbers increase significantly. Three upregulated srRNAs (OmrB, an outer member associated srRNA; RdlB, a swarming motility and curli expression regulator; McaS, a flagellar motility and biofilm formation regulator) overlapping in S2/S1 and S3/S1, together with the other four upregulated srRNAs (MicF, a ribosome binding inhibitor; MicL, an outer membrane associated srRNA; RybB, a cell envelope stress regulator; RydB, regulator of a global regulator RpoS) in S2/S1 are of special interest. By constructing deletion mutants and overexpression strains in uropathogenic *E. coli* strain UTI89, we tested their persister-formation capabilities in log phase and stationary phase cultures exposed to antibiotics (gentamicin, cefotaxime and levofloxacin) and stresses (heat, hyperosmosis, H_2_O_2_, and acid). The results of the deletion mutant studies showed that all the seven identified sRNAs have varying effects on persister formation with different antibiotics or stresses. Moreover, we found all the deletion mutants of these srRNAs have reduced biofilm formation. Additionally, except the McaS and the RydB overexpression strains, all of the srRNAs overexpression strains demonstrated increased persister-formation in antibiotic and stress persister assays, confirming the role of these srRNAs in persistence. Together, we identified seven srRNAs (OmrB, RdlB, McaS, MicF, MicL, RybB, and RydB) that are involved in type II persister formation for the first time. These findings provide convincing evidence for a new level of rapid persistence regulation via srRNA and furnish novel therapeutic targets for intervention.

## Introduction

Persisters have drawn wide attention since they can be identified in almost every bacterial species, even in fungi and eukaryotic human cancer cells [1-5]. They refer to a small subpopulation of dormant cells that can survive lethal antibiotics or stresses and regain susceptibilities when regrowing in fresh medium [6-8]. Persisters account for the recalcitrance of treatment of many persistent bacterial infections and can facilitate the emergence of antibiotic resistant bacteria [7-11]. Consequently, persisters pose great challenges for effective treatment of many bacterial infections

Persisters are divided into, type I and type II persisters [12]. Type I persisters stem from the stationary phase, while type II persisters are induced by triggering environmental signals as the cultures grow older from log phase to stationary phase. Unlike type I persister, type II persisters make up a great majority of the persisters in the stationary phase [13] and we consider them very important for understanding persister-formation mechanisms. Although various persister genes have been identified, how persisters are formed from a growing cell to a persister cell is still unclear.

Small regulatory RNAs (srRNAs) are widely spread in three domains of life, Archaea, Bacteria, and Eukaryotes [14]. In *Escherichia coli (E. coli),* they are non-coding, 50-500 nucleotides in length and synthesized under specific conditions [15-17]. srRNAs function in stress response, virulence regulation, biofilm formation, cell motility, uptake and metabolisms [18-22]. Though various functions have been described, their roles in persistence remain unclear. It is critical to determine the persister-formation capacities of srRNAs as they can respond to external signals quickly without protein synthesis.

To study the role of srRNA in persistence, especially those related with the emergence of type II persisters, we systematically interrogated the expression levels of the known srRNAs at three important timepoints where persisters switch from zero to high numbers. By constructing deletion mutants and overexpression strains of the candidate srRNAs and challenging them with different antibiotics and stresses, we identified seven srRNAs which play critical roles in regulating persister-formation.

## Materials and methods

### Bacterial strains, plasmids, and growth conditions

*E. coli* K12 strain W3110 and uropathogenic *E. coli* strain UTI89 as well as its derivatives were used in our experiments. The plasmid pBAD202 was used to construct the overexpression strains. The medium was supplemented with kanamycin (50 μg/mL) or chloramphenicol (25 μg/mL) to maintain resistance where necessary. Bacteria stored in -80°C were transferred to fresh Luria-Bertani (LB) broth (10 g Bacto-tryptone, 5 g yeast extract, and 10 g NaCl/liter) at 37°C, 200 rpm and grew overnight before use. Bacteria were diluted 1:1000 and routinely regrown in LB broth at 37°C, 200 rpm in our experiments, unless otherwise stated.

### Construction of deletion mutants and overexpression strains

sRNA deletion mutants were constructed successfully using the λ Red recombination system as described by Datsenko and Wanner [23]. Primers used to amplify all knockout-DNA fragments and to verify the correct constructs by polymerase chain reaction (PCR) are shown in Tables S1 and S2.

The arabinose-inducible plasmid pBAD202 was used to construct overexpression strains [24]. The primers used for the construction of the plasmid are listed in Table S3. Genes were amplified with primers and digested with the restriction enzymes *NcoI* and EcoRI (New England Biolabs). pBAD202 was also digested with the two enzymes and used for ligation with the PCR fragments using the T4 DNA ligase (New England Biolabs). The new constructs along with the empty vector, pBAD202, were transformed into parent strain UTI89 for overexpression experiments. The deletion mutants and overexpression strains were verified by DNA sequencing. Arabinose (0.1%) was added to the cultures of overexpression strains to induce the conditional expression of candidate genes [3].

### RNA isolation and quantitative real time-PCR (RT-PCR)

Bacteria were routinely cultured in LB medium followed by centrifugation at 4°C, 5000 rpm to remove the supernatant. RNAprotect Bacteria Reagent (Qiagen) was added to resuspended cell suspensions immediately. Total RNA was isolated from cells using bacterial RNA kit (Omega Bio-tek) according to the manufacturer’s protocol. Standardized total RNA was converted to cDNA using PrimeScript TMRT reagent Kit with gDNA Eraser (Takara) as described by the manufacturer. cDNA was then used as template to perform RT-PCR on an Applied Biosystems 7500 real-time instrument. The 16S rRNA gene *rrsB* was used as the reference gene.

### Persister assay

Persister levels were determined by counting the number of colony forming units (CFUs) that grew on LB agar plates without antibiotics following exposure to antibiotics, washing, and serial dilutions as previously described [3]. The antibiotics levofloxacin (5 μg/mL), cefotaxime (128 μg/mL), gentamicin (30 μg/mL) were added directly to cultures at the exponential phase (3 h of cultivation, ~10^8^ CFU/ mL) or stationary phase (10 h of cultivation, ~10^9^ CFU/ mL). Aliquots of the bacterial cultures were incubated at 37°C at different time points. To determine CFU, cultures were washed in phosphate-buffered saline before plating on LB plates in the absence of antibiotics [25].

### Susceptibility to other stresses in exposure assays

For acid stress (pH 3.0) and hyperosmosis stress (NaCl, 4 M), cultures were washed twice and resuspended in the same volume of the corresponding LB medium (pH 3.0, adjusted with HCl or NaCl, 4 M, respectively). For heat shock, bacteria were put in a water bath at 53°C for 1 h or 2 h. For the oxidative stress test, stationary phase cultures were diluted 1:100 with LB and exposed to hydrogen peroxide (H_2_O_2_) at a final concentration of 10 mM for up to 30 minutes. After exposure to various stresses, bacteria were washed in phosphate-buffered saline before plating on LB plates in the absence of antibiotics to determine CFU count [25, 26].

### Biofilm assay

Biofilm assays were performed as described previously [27]. Overnight cultures grown in LB were diluted to OD_600_ of ~0.05 in LB. A 200 μl aliquot of the diluted culture was added to each well of a 96-well polystyrene microtiter plate (Sermo, USA) and inoculated at 37°C without shaking for 24 h. The planktonic cells were determined by measuring OD_600_ using the SpectraMax Paradigm multi-mode detection platform (Applied Biosystems). The plate was washed with distilled water to remove the planktonic cells and retained with 220 μl of 0.1% crystal violet for 10 min. Unattached dye was rinsed away by washing with water for three times. Then the plate was dried and added with acetic acid (30%) to solubilize fixed crystal violet. The fixed biofilms were detected at OD_570_ and normalized by OD_600_.

### Statistical analysis

All experiments were performed at least in triplicate. The Mann-Whitney *U* test (non – parametric tests) in Prism 6.0 software (GraphPad, La Jolla, CA, USA) was used to analyze the data (UTI89 vs. mutants) to determine the statistical significance of differences [28]. Error bars indicated standard deviations, and all data were presented as the mean ± standard deviation. A *P* < 0.05 was considered statistically significant.

## Results

### Determination of three timepoints for type II persisters upon ampicillin treatment

Type II persisters are induced by fluctuating environmental signals, not originated from passage through the stationary phase [12]. Thus, capturing the timepoints that type II persisters appear and increase significantly is very important to study the genes involved in type II persister formation. In order to minimize interference with type I persisters, *E. coli* strain W3110 overnight cultures were diluted to a very high dilution 1:10^5^ into fresh LB medium and then incubated at 37°C, with shaking at 100 rpm. Aliquots of bacteria were removed and exposed to ampicillin (100 μg/mL) for 3 hours to determine the persister numbers. We found that no persisters existed in the first three hours. However, persisters started to appear at the 4^th^ hour, with 2 ~3 CFU/mL being detected. At the 5^th^ hour, the persister number increased to ~ 42 CFU/mL (Fig. 1A). Meanwhile, the initial cell numbers at the 3^rd^, 4^th^ and 5^th^ hours were also counted, with bacterial concentration reaching ~2.1×10^5^, ~1.4×10^6^, ~1.1×10^7^ respectively (Fig. 1B). Results indicated that cell density is associated with the emergence of persisters. We referred to the 3^rd^ hour as stage 1 (S1) where no persisters were present at this time point. The 4^th^ hour is referred to as stage 2 (S2) where persisters were just beginning to appear, and the 5^th^ hour is referred to as stage 3 (S3) where persister numbers increased significantly at this timepoint. S1, S2 and S3 are three important timepoints we used to identify genes associated with type II persister formation (see below).

**Fig. 1.**
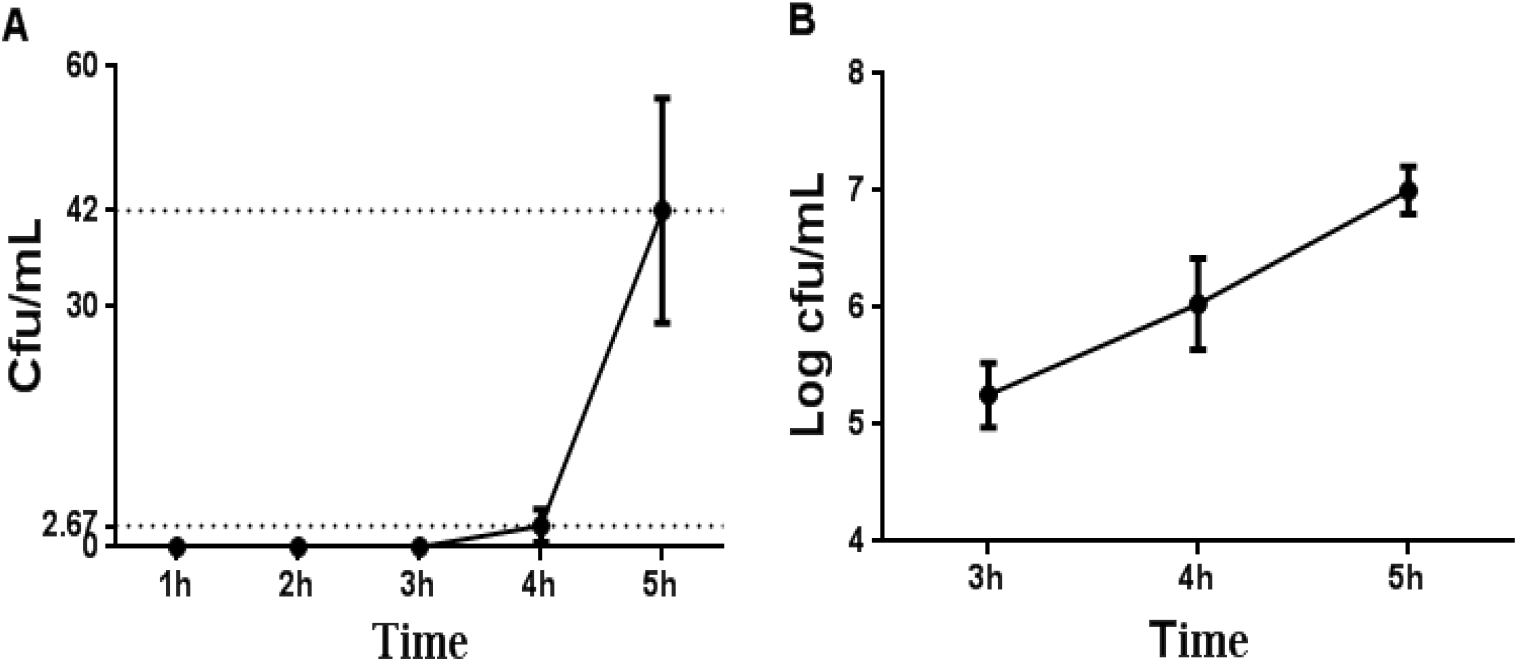
Determination of three timepoints for persister formation upon ampicillin treatment. (A) Persister numbers at each hour. Overnight *E. coli* K-12 W3110 was 1: 10^5^ diluted into fresh LB medium and then incubated at 37°C, 100 rpm. At each hour, aliquots were taken and exposed to ampicillin (100 μg/mL) for 3 hours to determine persister numbers. (B) The initial cell count. At the indicated hour, aliquots were taken without any treatment to determine the cell numbers directly. The experiments were performed in triplicate, and error bars represent standard deviations.

### Different expression levels of small regulatory RNAs in S1, S2, S3

To study the expression levels of srRNAs at the timepoints when persisters first appeared and then increased, we determined the expression levels of all the 57 srRNAs in the comprehensive EcoCyc database [29]. We found eight upregulated srRNAs (MicF, MgrR, MicL, OmrB, RdlB, McaS, RybB, RydB) (Fig. 2A) and 16 downregulated srRNAs (DicF, GadY, GlmZ, GlmY, IstR-1, OxyS, SgrS, SymR, SsrS, ArrS, FnrS, Och5, OhsC, RyfA, RyfD, RyeB) when comparing expression levels of S2 with S1. Six upregulated srRNAs (OmrB, RseX, McaS, MgrR, MicA, RdlB) (Fig. 2B) and 17 downregulated srRNAs (GadY, GlmZ, GlmY, IstR-1, OmrA, OxyS, RdlD, RybB, SgrS, SsrS, FnrS, Och5, RyfA, RyfD, RyeB, SibD, SibE) were detected when comparing expression levels of S3 with S1. Similarly, three upregulated srRNAs (DicF, ArrS, RyeG) and seven downregulated srRNAs (DsrA, GadY, RdlD, RybB, RydB, RyfA, SibE) were observed when comparing expression levels of S3 with S2. Target genes with known functions of the upregulated srRNAs are shown in Table 1. Pathways of these target genes comprise energy production, transport system, TA module, biofilm formation, global regulator, protease, trans-translation system, efflux pump that belong to known persister pathways [7], indicating the upregulated srRNAs might participate in persister formation.

**Fig. 2.**
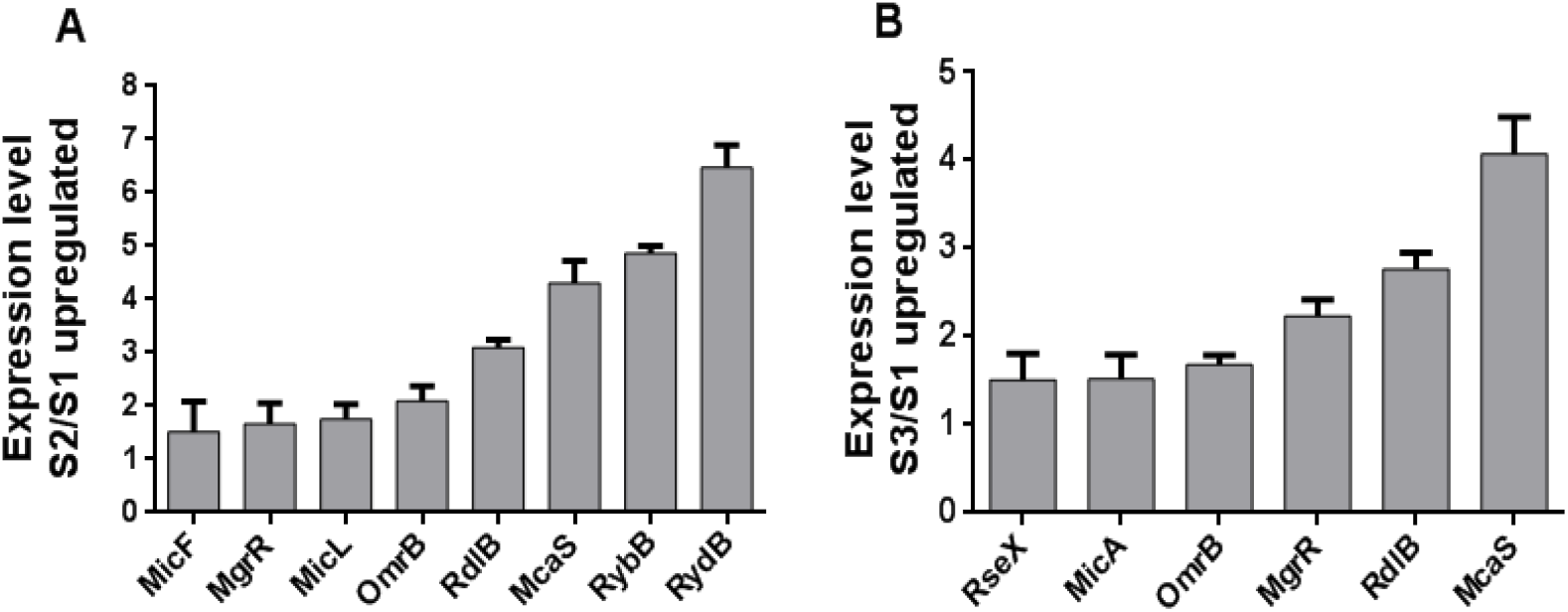
Upregulated small regulatory RNAs in S2/S1 and S3/S1. Overnight *E. coli* K-12 W3110 was 1:10^5^ diluted into fresh LB medium and then incubated at 37°C, 100 rpm up to S1 (3 hours), S2 (4 hours) and S3 (5 hours), respectively. And then, total RNA was extracted and used for qRT-PCR. Upregulated small regulatory RNAs when comparing srRNA expression levels in S2/S1 (A), S3/S1 (B) are shown. The error bars show standard deviations (n=3).

**Table 1.**
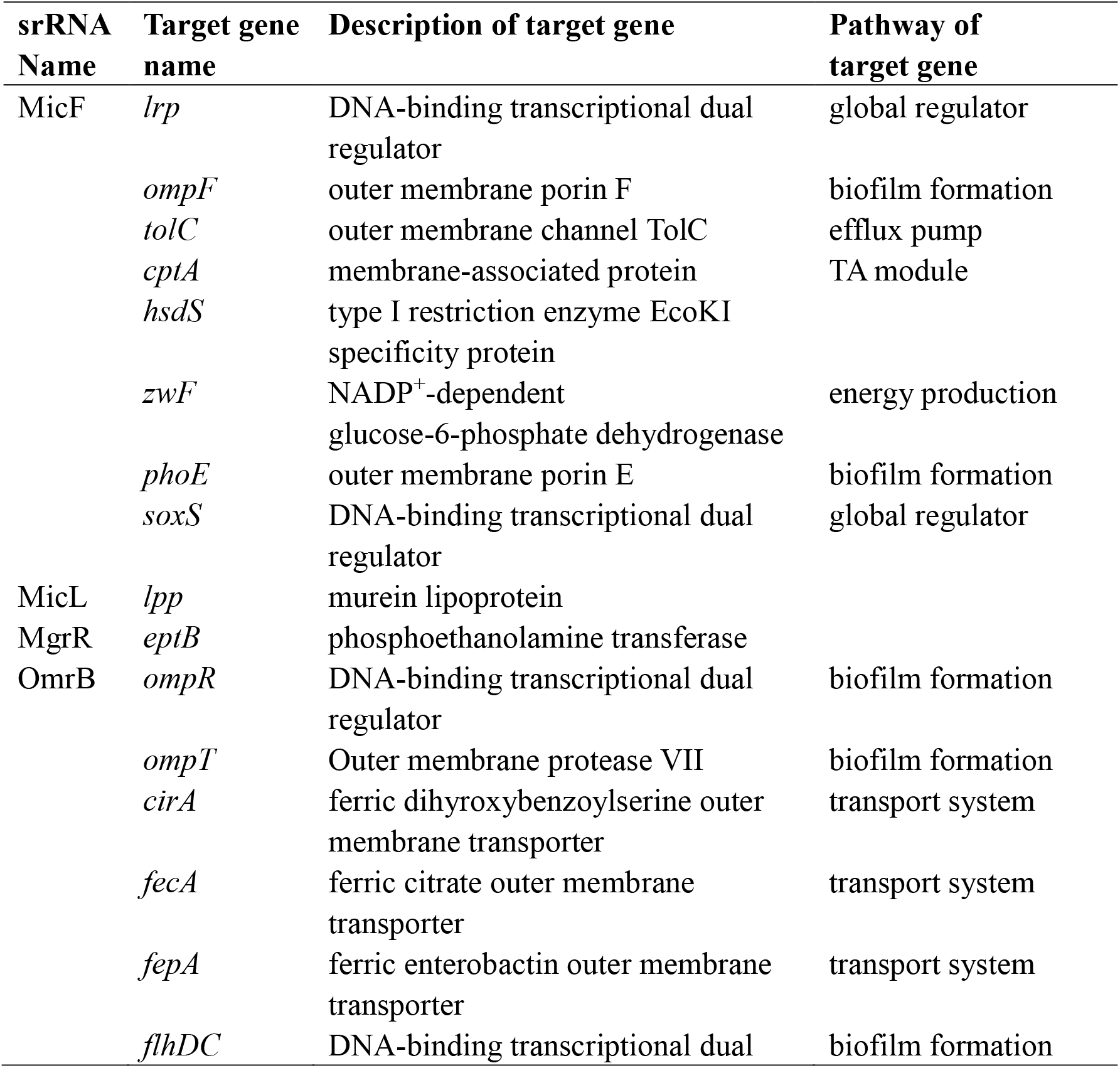
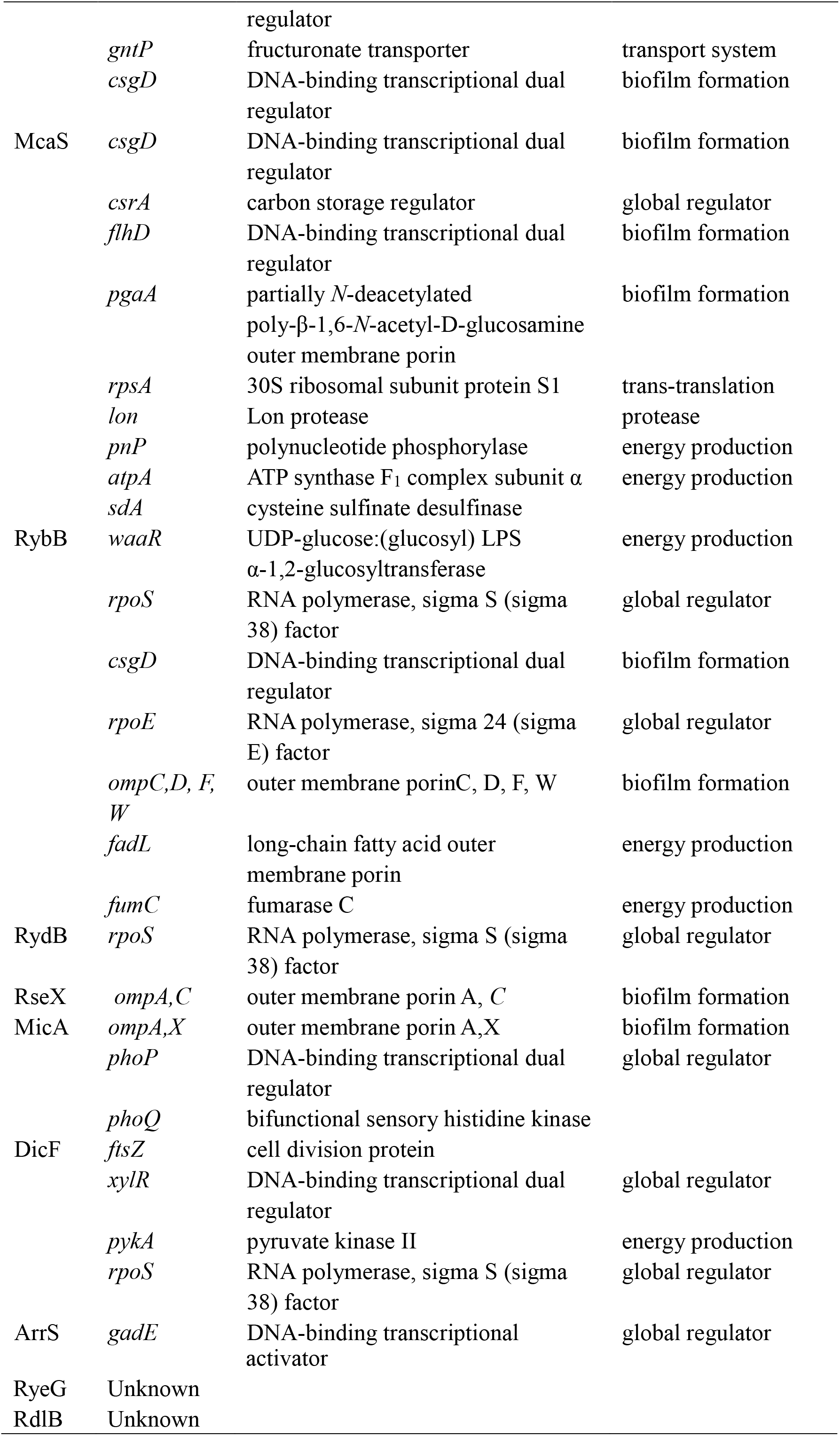
Target genes of the upregulated srRNAs.

| srRNA Name | Target gene name | Description of target gene | Pathway of target gene |
| --- | --- | --- | --- |
| MicF | *lrp* | DNA-binding transcriptional dual regulator | global regulator |
|  | *ompF* | outer membrane porin F | biofilm formation |
|  | *tolC* | outer membrane channel TolC | efflux pump |
|  | *cptA* | membrane-associated protein | TA module |
|  | *hsdS* | type I restriction enzyme EcoKI specificity protein |  |
|  | *zwF* | NADP<sup>+</sup>-dependent glucose-6-phosphate dehydrogenase | energy production |
|  | *phoE* | outer membrane porin E | biofilm formation |
|  | *soxS* | DNA-binding transcriptional dual regulator | global regulator |
| MicL | *lpp* | murein lipoprotein |  |
| MgrR | *eptB* | phosphoethanolamine transferase |  |
| OmrB | *ompR* | DNA-binding transcriptional dual regulator | biofilm formation |
|  | *ompT* | Outer membrane protease VII | biofilm formation |
|  | *cirA* | ferric dihydroxybenzoylserine outer membrane transporter | transport system |
|  | *fecA* | ferric citrate outer membrane transporter | transport system |
|  | *fepA* | ferric enterobactin outer membrane transporter | transport system |
|  | *flhDC* | DNA-binding transcriptional dual regulator | biofilm formation |
|  | *gntP* | fructuronate transporter | transport system |
|  | *csgD* | DNA-binding transcriptional dual regulator | biofilm formation |
| McaS | *csgD* | DNA-binding transcriptional dual regulator | biofilm formation |
|  | *csrA* | carbon storage regulator | global regulator |
|  | *flhD* | DNA-binding transcriptional dual regulator | biofilm formation |
| | *pgaA* | partially *N*-deacetylated poly-β-1,6-*N*-acetyl-D-glucosamine outer membrane porin | biofilm formation |
|  | *rpsA* | 30S ribosomal subunit protein S1 | trans-translation |
|  | *lon* | Lon protease | protease |
|  | *pnP* | polynucleotide phosphorylase | energy production |
| | *atpA* | ATP synthase F<sub>1</sub> complex subunit $\alpha$ | energy production |
|  | *sda* | cysteine sulfinase desulfinate |  |
| RybB | *waaR* | UDP-glucose:(glucosyl) LPS α-1,2-glucosyltransferase | energy production |
|  | *rpoS* | RNA polymerase, sigma S (sigma 38) factor | global regulator |
|  | *csgD* | DNA-binding transcriptional dual regulator | biofilm formation |
|  | *rpoE* | RNA polymerase, sigma 24 (sigma E) factor | global regulator |
|  | *ompC,D, F, W* | outer membrane porinC, D, F, W | biofilm formation |
|  | *fadL* | long-chain fatty acid outer membrane porin | energy production |
|  | *fumC* | fumarase C | energy production |
| RydB | *rpoS* | RNA polymerase, sigma S (sigma 38) factor | global regulator |
| RseX | *ompA,C* | outer membrane porin A, C | biofilm formation |
| MicA | *ompA,X* | outer membrane porin A,X | biofilm formation |
|  | *phoP* | DNA-binding transcriptional dual regulator | global regulator |
|  | *phoQ* | bifunctional sensory histidine kinase |  |
| DicF | *ftsZ* | cell division protein |  |
|  | *xylR* | DNA-binding transcriptional dual regulator | global regulator |
|  | *pykA* | pyruvate kinase II | energy production |
|  | *rpoS* | RNA polymerase, sigma S (sigma 38) factor | global regulator |
| ArrS | *gadE* | DNA-binding transcriptional | global regulator |

|  |  |  |
| --- | --- | --- |
|  |  | activator |
| RyeG | Unknown |  |
| RdlB | Unknown |  |

### Susceptibility of the srRNA deletion mutants to different antibiotics

To determine the role of the seven upregulated srRNAs in persistence, we constructed their deletion mutants *(ΔmicF, ΔomrB, ΔrybB, ΔmcaS, ΔmicL, ΔrdlB, ΔrydB)* in the uropathogenic *E. coli* strain UTI89 and tested their persister phenotypes to different classes of antibiotics. We monitored the numbers of the parent strain and all the deletion mutants without any antibiotic or stress exposure from the beginning to the end of the experiments and no difference was observed between the parent strain and the mutant strains (data not shown). In the log phase cultures (~10^8^ CFU/mL), when all the bacteria were challenged with gentamicin (30 μg/mL), Δ*mcaS, ΔrydB* and Δ*micL* mutants showed decreased persister numbers, ranging from 7-fold *(ΔmicL)* to 10-fold (Δ *mcaS)* difference compared with the parent strain, whereas the other mutants were not affected (Fig. 3A). When exposed to cefotaxime (128 μg/mL), all the mutants showed a decrease in persistence compared with the parent strain UTI89, ranging from 8-fold *(ΔrdlB)* to 170-fold *(ΔmcaS)* (Fig. 3B). The log phase cultures were also treated with levofloxacin for 0.5 h, and there was about 5-log decrease in all the tested strains, with no difference observed between the deletion mutants and the parent strain (data not shown).

**Fig. 3.**
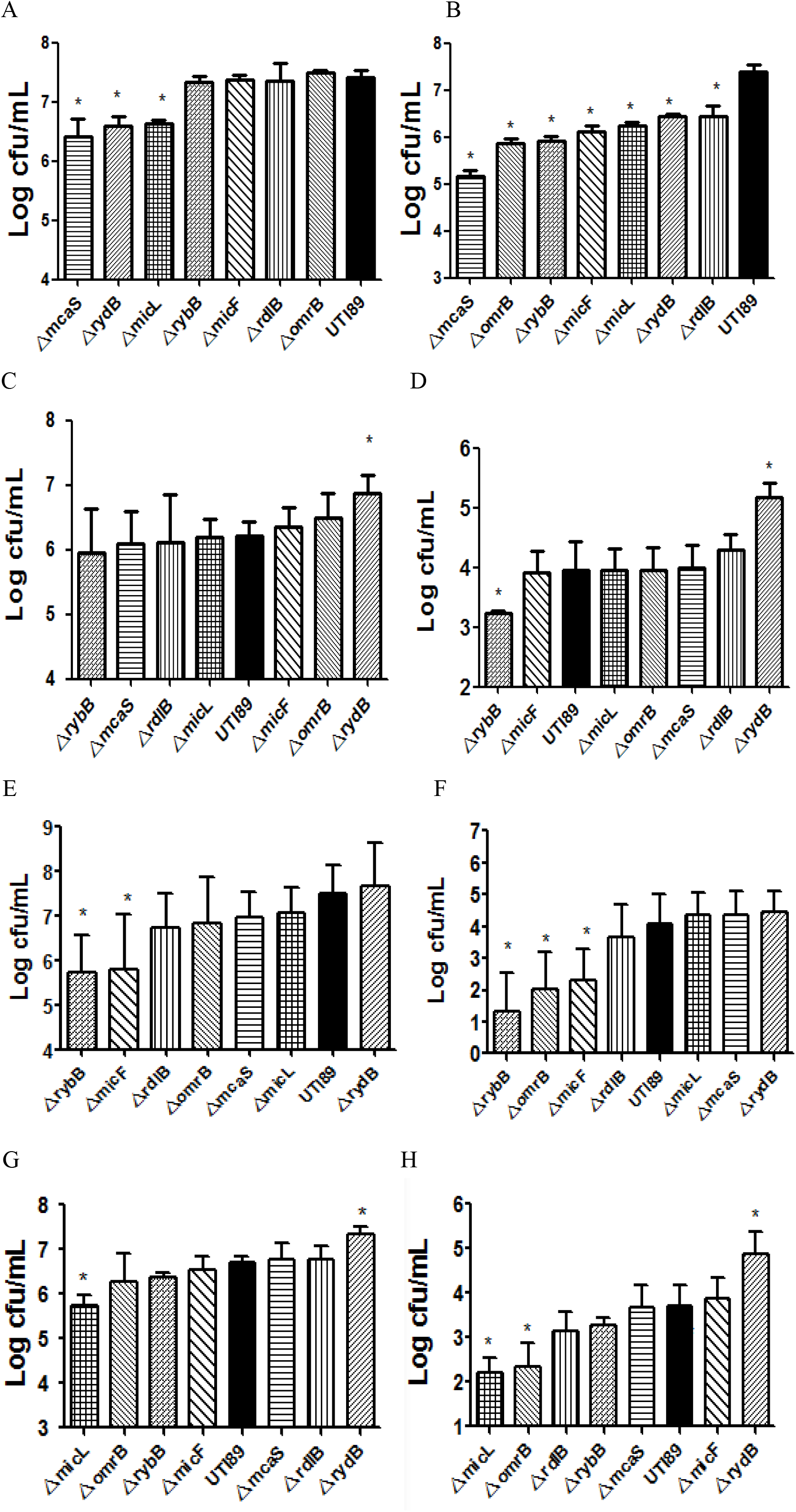
Susceptibility of the srRNA deletion mutants to antibiotics. (A, B) Susceptibilities of the log phase bacteria to gentamicin (30 μg/mL, A) or cefotaxime (128 μg/Ml, B) after exposure for six hours. (C, D) Susceptibilities of the stationary phase bacteria to levofloxacin (5 μg/mL) on the second day (C) or the sixth day (D). (E, F) Susceptibilities of the stationary phase bacteria to gentamicin (30 μg/mL) on the second day (E) or the third day (F). (G, H) Susceptibilities of the stationary phase bacteria to cefotaxime (128 μg/mL) on the forth day (G) or the sixth day (H). Bacteria were cultured to log phase (~10^8^ CFU/mL) or stationary phase (~10^9^ CFU/mL) prior to the addition of antibiotics. The number of persisters was determined at the time point indicated in the figure. The error bars show standard deviations (n=3). * Indicates significant difference from the other strains at *p* < 0.05.

However, when the stationary phase cultures were exposed to levofloxacin for two days, the persister phenotype could be observed. Δ*rydB* mutant showed a dramatic increase in persistence (Fig. 3C). After exposure to levofloxacin for six days, it could be observed that another deletion mutant *(ΔrybB)* had a decreased persister phenotype (Fig. 3D), indicating *rybB* could be a late persister gene. When the seven srRNAs deletion mutants were treated with gentamicin for two days, Δ*rybB* and Δ*micF* mutants had prominent decrease in persistence compared to the parent strain UTI89 (Fig. 3E), while the Δ*omrB* mutant demonstrated persister defect at day 3 (Fig. 3F). After exposure to cefotaxime, the Δ*micL* mutant showed defect in persister formation, whereas the Δ*rydB* mutant showed increased persister-formation level at day 4 (Fig. 3G). The Δ*omrB* mutant also had lower persister number upon cefotaxime treatment at day 6 (Fig. 3H).

### Susceptibility of the srRNA deletion mutants to stresses

To determine the effect of stresses on the survival of the srRNA deletion mutants, we subjected all the seven mutants to hyperosmosis (NaCl, 4M), acid (pH 3.0), heat (53°C) and oxidative (H_2_O_2_, 10 mM) stresses and assessed their survival in the stationary phase (Fig. 4). Upon exposure to hyperosmosis for five days, only the Δ*micL* mutant showed dramatic decrease in survival (208-fold decrease) compared with the parent strain UTI89. No difference could be observed between the other deletion mutant strains and the parent strain (Fig. 4A). When treated with acid, four deletion mutants *(ΔmicL, ΔomrB, ΔrdlB, ΔrybB)* showed higher susceptibilities (at least 16-fold decrease) compared with the parent strain, while the other three deletion mutants *(ΔmicF, ΔmcaS, ΔrydB)* had similar susceptibilities as the parent strain (Fig. 4B). Upon exposure to heat, five deletion mutants *(ΔmicL, ΔomrB, ΔmicF, ΔrdlB, ΔrybB)* showed higher susceptibilities (at least 13-fold decrease) when compared with the parent strain (Fig. 4C). When all the deletion mutants were subjected to H_2_O_2_ oxidative stress, the survival of the five deletion mutant strains *(ΔrdlB, ΔmicL, ΔrydB, ΔmicF, ΔmcaS)* was significantly decreased, whereas the other two mutants *(ΔomrB, ΔrybB)* showed the same magnitude of decrease as the parent strain (Fig. 4D).

**Fig. 4.**
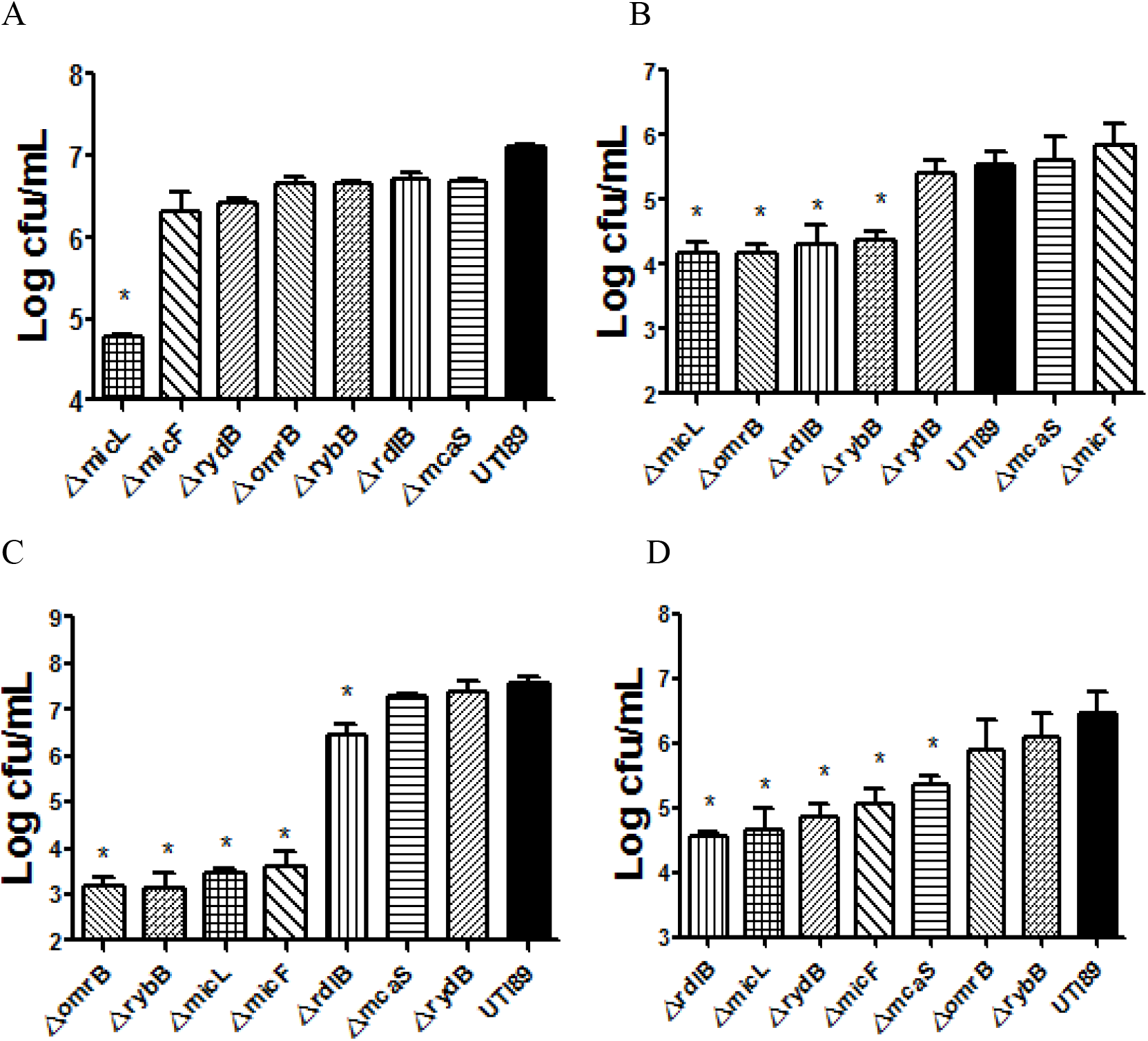
Susceptibility of the srRNA deletion mutants to stresses. (A) Susceptibilities to hyperosmosis (NaCl, 4 M) after exposure for five days. (B) Susceptibilities to acid (pH 3.0) after exposure for one day. (C) Susceptibilities to heat (53°C) after exposure for two hours. (D) Susceptibilities to oxidative stress (H_2_O_2_, 10 mM) after exposure for 0.5 hour. Bacteria were cultured to the stationary phase (~10^9^ CFU/mL) prior to the addition of stresses. Detailed methods were described above. The error bars show standard deviations (n=3). * Indicates significant difference from the other strains at *p* < 0.05.

### Effects of the overexpression of srRNAs on persister levels

The seven srRNAs *(ΔmicF, ΔomrB, ΔrybB, ΔmcaS, ΔmicL, ΔrdlB, ΔrydB)* were overexpressed to further characterize their effects on persistence. The newly constructed plasmids carrying the corresponding srRNAs along with the empty vector pBAD202 were transformed into parent strain UTI89 for construction of the overexpression strains. In the stationary phase cultures, we found the five srRNAs (MicF, MicL, OmrB, RdlB, RybB) overexpression strains showed higher persister levels than the control strain upon exposure to levofloxacin (5μg /mL), gentamicin (30 μg /mL), cefotaxime (128 μg /mL) and various stresses hyperosmosis (NaCl, 4M), acid (pH, 3.0), heat (53°C), and oxidative stress (H_2_O_2_, 10 mM), respectively. However, the RydB overexpression strain resulted in decreased persister levels to levofloxacin, cefotaxime, hyperosmosis and heat exposure, but showed increased persister levels to gentamicin, acid and oxidation treatment. The McaS overexpression strain also demonstrated higher persister-formation capabilities to levofloxacin, cefotaxime, gentamicin and stresses hyperosmosis, acid and oxidation exposure, whereas it had defect when challenged with heat stress (Table 2-3). Overall, all of the seven srRNAs have varying effects on persister-formation when overexpressed.

**Table 2.** Effect of overexpression of srRNAs on persister levels upon antibiotic exposure. Stationary phase bacteria (~10^9^ CFU/mL) were challenged with levofloxacin (LVX, 5 μg /mL), gentamicin (GEN, 30 μg/mL) and cefotaxime (CTX, 128 μg/mL), respectively, for five days. CFU values were determined at the appropriate times.

| Overexpression strains | No. of bacteria (Log CFU/mL) |  |  |  |
| --- | --- | --- | --- | --- |
|  | Starting CFU | LVX | GEN | CTX |
| UTI89-Pvector | 9.4±0.12 | 3.3±0.21 | 2.5±0.45 | 6.1±0.09 |
| UTI89-PmcaS | 9.8±0.05 | 5.3±0.07 | 4.2±0.13 | 7.9±0.13 |
| UTI89-PrdlB | 9.7±0.05 | 6.0±0.02 | 4.6±0.08 | 8.0±0.10 |
| UTI89-PrydB | 9.7±0.04 | 2.2±0.18 | 5.3±1.55 | 0 |
| UTI89-PomrB | 9.8±0.02 | 6.1±0.06 | 3.9±0.28 | 8.1±0.02 |
| UTI89-PmicL | 9.8±0.02 | 5.4±0.63 | 4.0±0.42 | 7.8±0.03 |
| UTI89-PrybB | 9.8±0.03 | 5.0±0.14 | 4.2±0.09 | 7.6±0.06 |
| UTI89-PmicF | 9.8±0.03 | 5.1±0.13 | 5.2±0.12 | 8.0±0.04 |

**Table 3.** Effect of overexpression of srRNAs on persister levels upon stress exposure. Stationary phase bacteria (~10^9^ CFU/mL) were directly treated with various stresses. For hyperosmosis or acid pH stress, the stationary phase bacteria were harvested, washed and then resuspended in hyperosmotic LB medium (NaCl, 4M) for three days as well as in acid LB medium (pH, 3.0) for one day. For heat, bacteria were directly put in a water bath of 53°C for one hour. For oxidative stress, bacteria were 1:100 diluted before exposure to 10 mM H_2_O_2_ for 0.5 hour. CFU values were determined at the appropriate times.

| Overexpression strains | No. of bacteria (Log CFU/mL) |  |  |  |  |
| --- | --- | --- | --- | --- | --- |
|  | Starting CFU | Hyper-osmosis | Acid | Heat | H<sub>2</sub>O<sub>2</sub> |
| UTI89-Pvector | 9.4±0.12 | 3.6±0.41 | 2.8±0.06 | 3.9±0.17 | 4.1±0.17 |
| UTI89-PmcaS | 9.8±0.05 | 7.8±0.40 | 4.0±0.08 | 2.0±0.25 | 5.9±0.22 |
| UTI89-PrdlB | 9.7±0.05 | 7.7±0.26 | 4.0±0.09 | 4.4±0.13 | 4.9±0.20 |
| UTI89-PrydB | 9.7±0.04 | 1.0±0.68 | 3.5±0.15 | 2.2±0.80 | 5.1±0.08 |
| UTI89-PomrB | 9.8±0.02 | 7.8±0.52 | 3.9±0.08 | 5.5±0.13 | 4.9±0.28 |
| UTI89-PmicL | 9.8±0.02 | 8.0±0.55 | 3.8±0.16 | 6.0±0.18 | 4.9±0.25 |
| UTI89-PrybB | 9.8±0.03 | 7.7±0.49 | 4.1±0.08 | 5.6±0.07 | 5.3±0.85 |
| UTI89-PmicF | 9.8±0.03 | 7.8±0.86 | 3.6±0.07 | 5.5±0.20 | 4.9±0.14 |

### Impact of the srRNA mutations on biofilm formation

Because the presence of persister cells in the biofilm contributes largely to the recalcitrance to antibiotic treatment [30-33], we determined the influence of the persister-associated srRNAs on biofilm-formation. Interestingly, the results showed all the deletion mutants *(ΔmicF, ΔomrB, ΔrybB, ΔmcaS, ΔmicL, ΔrdlB, ΔrydB)* had reduced biofilm formation to some degree compared with the parent strain UTI89 (Fig. 5). Among them, the Δ*mcaS* mutant was the weakest (~21% decrease) while the Δ*micF* mutant had the strongest effect on biofilm formation (~40% decrease) compared with the parent strain UTI89.

**Fig. 5.**
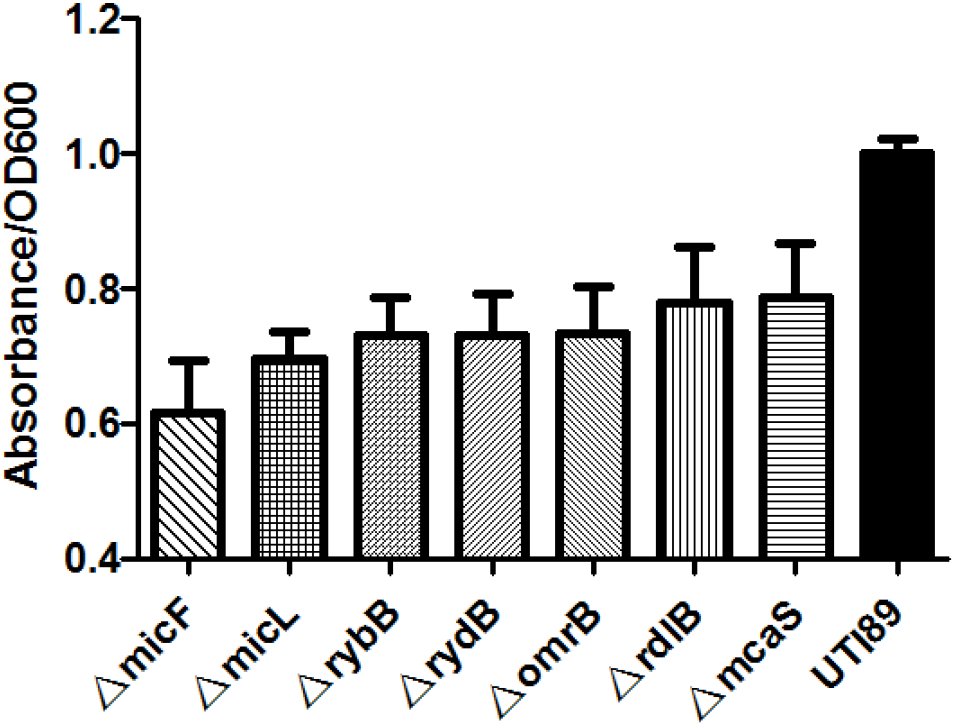
Effect of the deletion mutants on biofilm formation. 200 μl bacteria were transferred to 96-well microtiter plates and grown at 37°C for 24 h in LB medium. Planktonic cells were measured at OD_600_. After washing, staining with crystal violet, and dissolution, final biofilm formation was measured at OD_570_ and normalized to OD_600_.

## Discussion

In this study, we identified seven novel srRNAs (MicF, OmrB, RybB, McaS, MicL, RdlB, RydB) associated with persister formation in the uropathogenic *E. coli* strain UTI89. To our knowledge, this is the first study that comprehensively characterized the role of srRNA in persister formation. By comparing different expression levels of srRNAs in the three important timepoints S1, S2 and S3 relevant to persister formation, we identified initial natural persister formation related srRNAs (MicF, OmrB, RybB, McaS, MicL, RdlB, RydB) in the absence of antibiotic background. Because persisters are produced by a stochastic process [8, 34], monitoring the expression levels of genes after antibiotic treatment to search for persister-associated genes is not suitable, as genes could be induced by antibiotics thereby confounding the real important persister formation genes [35]. By using this new persister gene search methodology comparing the expression of srRNAs at different critical stage of no persister and persister emergence in our study, we were able to find seven native srRNAs (MicF, OmrB, RybB, McaS, MicL, RdlB, RydB) involved in type II persister formation.

Apart from their roles in exponential phase persister-formation (Fig. 3A, 3B), we found the seven srRNAs are also involved in persistence in stationary phase (Fig. 3C-3H, Fig. 4 and Table 2-3). This phenomenon suggests that srRNAs fulfill their persistence function in the whole life span of the bacteria and confirms that srRNAs play a vital role in persistence to different antibiotics and stresses.

McaS is a multi-cellular adhesive small regulatory RNA and is known to regulate flagellar motility and biofilm formation by targeting the global transcription regulators *(csgD, flhD)* and biofilm formation related mRNA *pgaA* [27, 36]. Protein co-purification study also found that McaS could bind with RpsA (translation initiation factor), Lon (DNA-binding ATP-dependent protease), PNPase (polynucleotide phosphorylase or - polymerase), AtpA (F1 sector of membrane-bound ATP synthase), CsrA (Regulatory McaS protein for carbon source metabolism) [37]. The targets of McaS match well to the known persister pathways including global regulator, biofilm formation, trans-translation, protease and energy production [7], which can count for the role of McaS in persistence. Notably, Lon protease can also be regulated by McaS. As we know, polyphosphated Lon protease can cause the degradation of antitoxins and the freed toxin, subsequently turning bacteria into dormant persisters [38, 39]. This suggests that McaS may also influence TA modules for persister formation.

MicF inhibits ribosome binding and induce the degradation of the RNA messages in many bacteria [40]. MicF can interact with *lrp* (a global regulator involved in amino acid biosynthesis and catabolism), and down-regulate translation of *ompF* (outer membrane porin), *soxS* (DNA-binding transcriptional dual regulator) and *tolC* (a central factor involved in efflux pump) [41], which can be important for persister formation and survival [41, 42]. By serving as a well known antisense RNA of *ompF,* MicF can down-regulate *ompF* to reduce the entry of antibiotics so persisters can survive. Pathways of these targets are involved in global regulator, biofilm formation and efflux pump activity. We propose that MicF may influence the persister pathways to cause persistence to antibiotics and stresses via its target genes. It has been shown different expression levels of MicF in response to osmolarity and temperature change [43], and our observation that MicF mutant has defective persistence to hyperosmosis and heat exposure is consistent with the previous study.

MicL was reported to have a sole target *lpp,* an abundant outer membrane lipoprotein in response to stress [44]. We suppose that MicL may influence *lpp* expression to affect transport of antibiotics and other stressors, thus facilitating persistence. In addition, using the updated tools CopraRNA and IntaRNA software for small RNA target prediction [45], we found MicL has binding sites with *mazF* (mRNA interferase toxin antitoxin) and *hipB* (HipB antitoxin / DNA-binding transcriptional repressor), which are two known persister genes involved in TA modules, indicating that MicL could mediate persister formation via these antitoxins.

RybB can regulate the expression of membrane porins related mRNA *(ompC, ompD, ompW, ompF,* and *fadL*) [46, 47], a transcriptional regulator associated with biofilm formation (csgD) [48], and also modulate LPS biosynthesis [49]. Additionally, overexpression of *rybB* causes increased expression of *rpoS* (a global regulator involved in persister formation) [50] and decreased levels of *fumC* (a fumarase isozyme participating in the TCA cycle) [46]. Thus, the targets of RybB match known persister pathways including those of biofilm formation, global regulators, and energy production [7], which could all account for persister phenotype of RybB.

The *omrB* deletion mutant showed defect in persistence when challenged with multiple antibiotics and stresses in both exponential phase and stationary phase cultures (Fig. 3-4). OmrB is known to regulate the expression levels of genes encoding many outer membrane proteins, including *cirA, fecA, fepA* and *ompT, ompR* [51], which could modulate outer membrane composition in response to environmental stress conditions. In addition, two transcriptional regulators *csgD* (important for biofilm formation) and *flhDC* together with a gluconate /fructuronate transporter *gntP* could also be modulated by OmrB [51-53]. Thus, we propose that OmrB can affect different persister pathways involved in biofilm, translational regulator and transport system to perform its persister function via its target genes.

To date, little is known about the functions of RdlB except that overexpression of RdlB decreases swarming motility and curli expression [54]. In this study, we demonstrated that RdlB is involved in persistence to antibiotics (levofloxacin, gentamicin, cefotaxime) and stresses (hyperosmosis, heat, oxidation, acid) for the first time (Fig. 3-4, Table 2-3). In order to investigate the mechanisms involved, we used the updated tools CopraRNA and IntaRNA for RdlB target prediction [45]. Results showed that RdlB has binding sites with energy production mRNA *srmB* (ATP-dependent RNA helicase), *paaH* (3-hydroxyadipyl-CoA dehydrogenase NAD+-dependent), transport system mRNA *nirC* (nitrite transporter), *ssuB* (aliphatic sulfonate ABC transporter ATPase), *potH* (putrescine ABC transporter permease), *nirD* (nitrite reductase NADH small subunit) and membrane associated mRNA *ybhM* (BAX Inhibitor-1 family inner membrane protein), *yiaT* (putative outer membrane protein), *pqiA* (inner membrane protein) as well as DNA repair mRNA *vsr* (DNA mismatch endonuclease of very short patch repair). The associated pathways of these targets are known to be associated with persistence [7]. Notably, RdlB could also bind to *tnaA,* a well-known persister gene connected to signaling pathway. Future studies are needed to confirm how RdlB is involved in persister formation.

So far, RydB is known to regulate *rpoS* (a global regulator involved in persister formation) expression only [55]. In our study, we found RydB could modulate persister formation when treated with antibiotics and lethal stresses. Using small RNA target prediction CopraRNA and IntaRNA software analysis [45], we found RydB could affect persister pathways [7] via energy production mRNA *sdhA* (succinate dehydrogenase flavoprotein subunit), *purC* (amidophosphoribosyltransferase), *yjfC* (ATP-Grasp family ATPase), *cbrA* (FAD-binding protein putative oxidoreductase), membrane associated mRNA *ybbJ* (inner membrane protein), *asmA* (inner membrane-anchored), *yihG* (inner membrane protein inner membrane acyltransferase), *yobD* (UPF0266 family inner membrane protein), transport system mRNA *yncD* (putative iron outer membrane transporter), *artQ* (arginine ABC transporter permease), *fecC* (ferric citrate ABC transporter permease), *satP* (succinate-acetate transporter), efflux pump mRNA *mdtC* (multidrug efflux system subunit C), *ydhK* (putative efflux protein, component of YdhJK efflux pump) and trans-translation system *rpsA* mRNA. Among them, *sdhA* and *rpsA* are known persister genes [56, 57].

Biofilm can resist antibiotic killing without any drug resistance mechanisms. The link between the presence of persisters and biofilm formation is the subject of many studies [31, 32, 58, 59]. We propose that the roles of the srRNAs in persistence are associated with their biofilm formation capability, based on our findings. In line with this proposition, we found that the deletion mutants of the seven srRNAs all showed defective biofilm formation (Fig. 5). In particular, our finding of McaS is consistent with the previous finding that McaS is associated with biofilm formation [27]. However, the impacts of srRNA on attachment and secretion of extracellular polymeric substances to the biofilm remains further study.

In summary, we identified seven new small regulatory RNAs (OmrB, RdlB, McaS, MicF, MicL, RybB, and RydB) that are involved in persister formation using a novel persister gene search methodology. The targets of these small regulatory RNAs involved in several persister pathways including energy production, transport system, SOS response, DNA repair, TA module, biofilm formation, global regulator, trans-translation system, and efflux pump. Future studies are needed to address the role of srRNAs in persister formation in other bacteria and *in vivo* as well as the exact mechanisms by which srRNAs regulate the persistence phenomenon. Our findings provide new insights into mechanisms of persister formation and its regulation at the srRNA level and offer new targets for treatment of persistent infections.

## Author Contributions

YZ, WZ designed the experiments and revised the manuscript; SZ and SL completed all the experiments, SZ and NW performed the data analysis; SZ and YZ wrote the manuscript.

## Acknowledgments

We thank Peng Cui, Tao Xu, Jing Wu and Jiazhen Chen, at Department of Infectious Diseases, Huashan Hospital, China for helpful advice in data analysis. The research was supported by the National Natural Science Foundation of China (81572046).

---

**Table S1.**
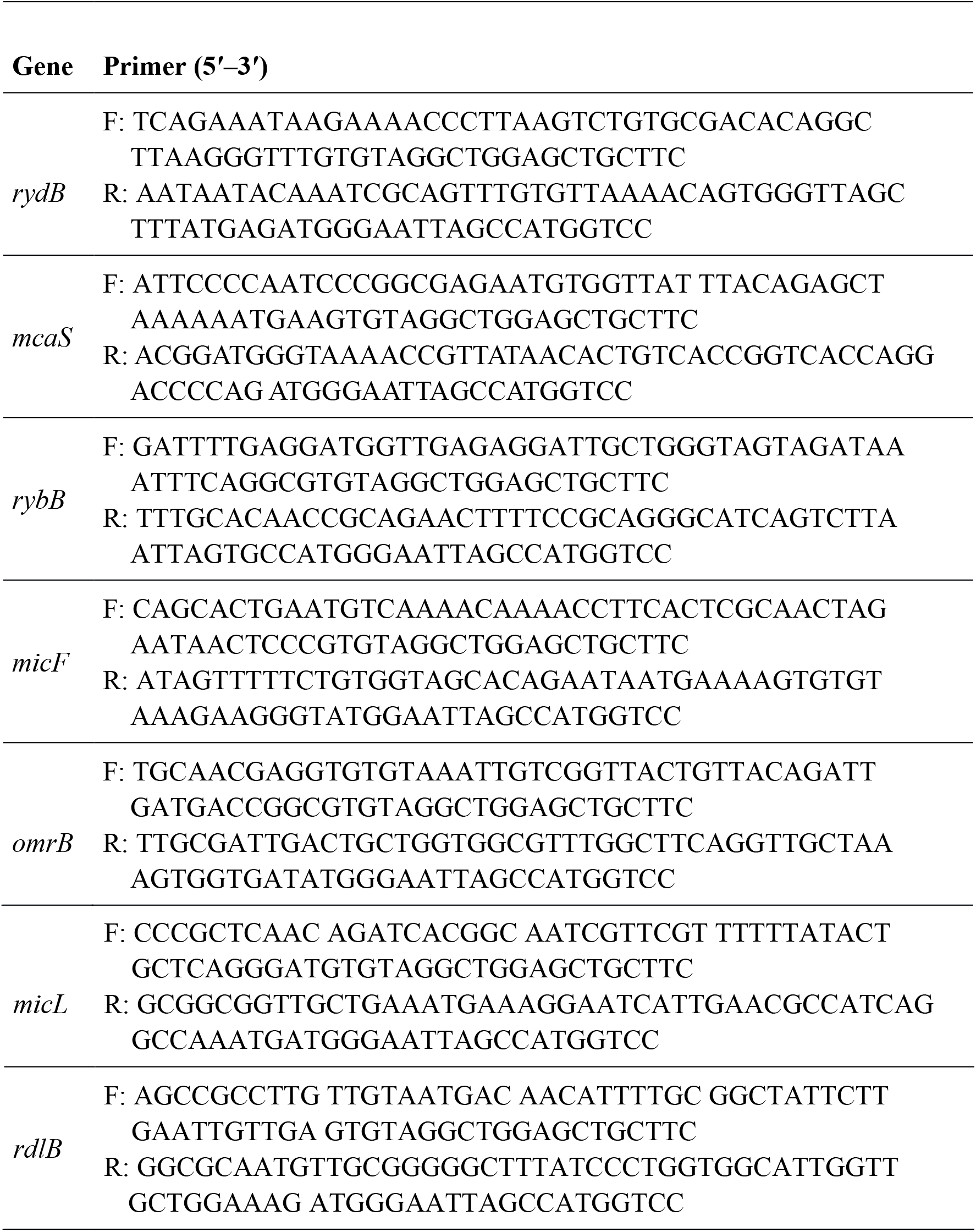
Primers used to generate gene-knockout mutants.

**Table S2.**
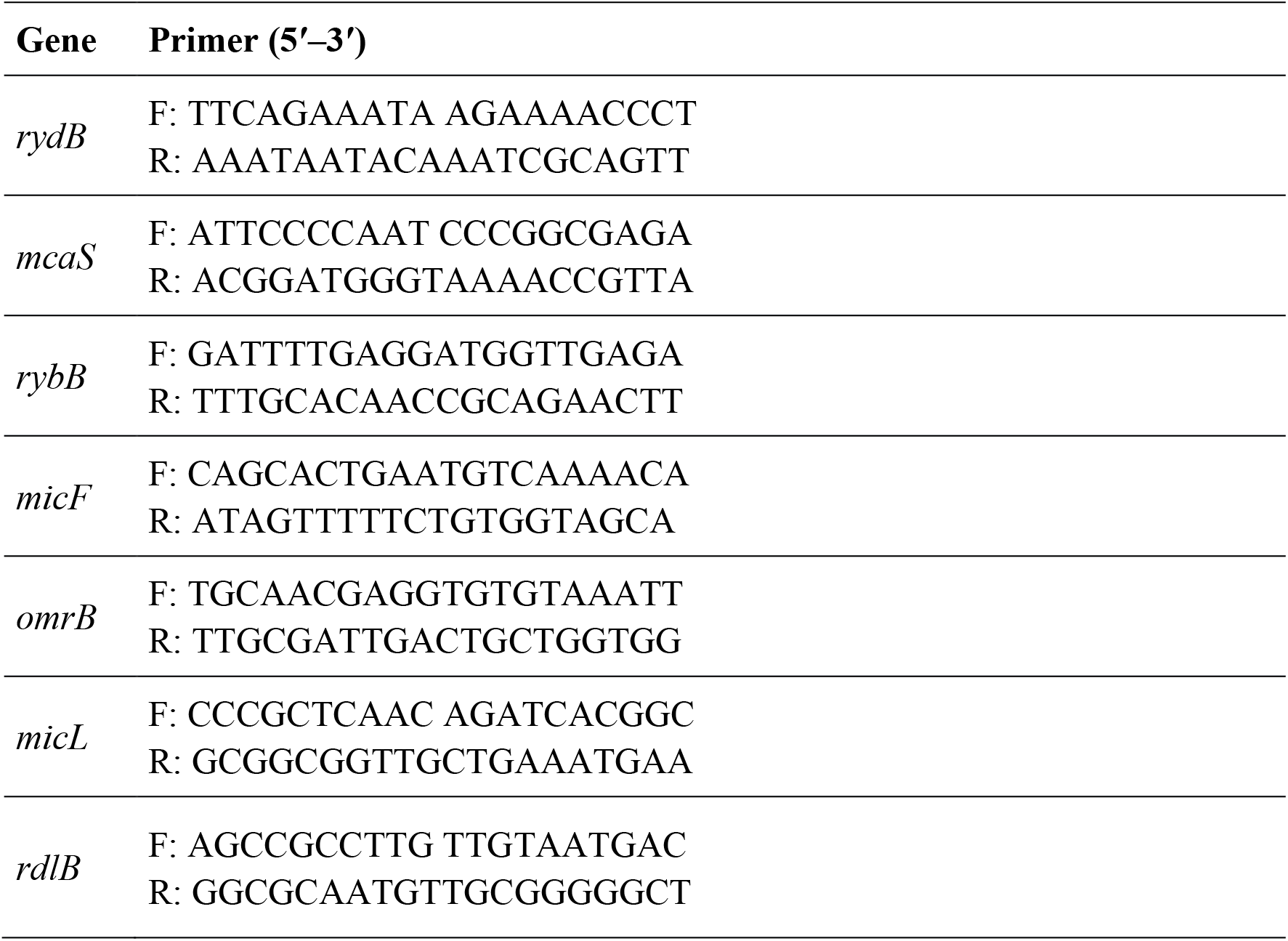
Primers used to verify gene knockout mutants.

**Table S3.**
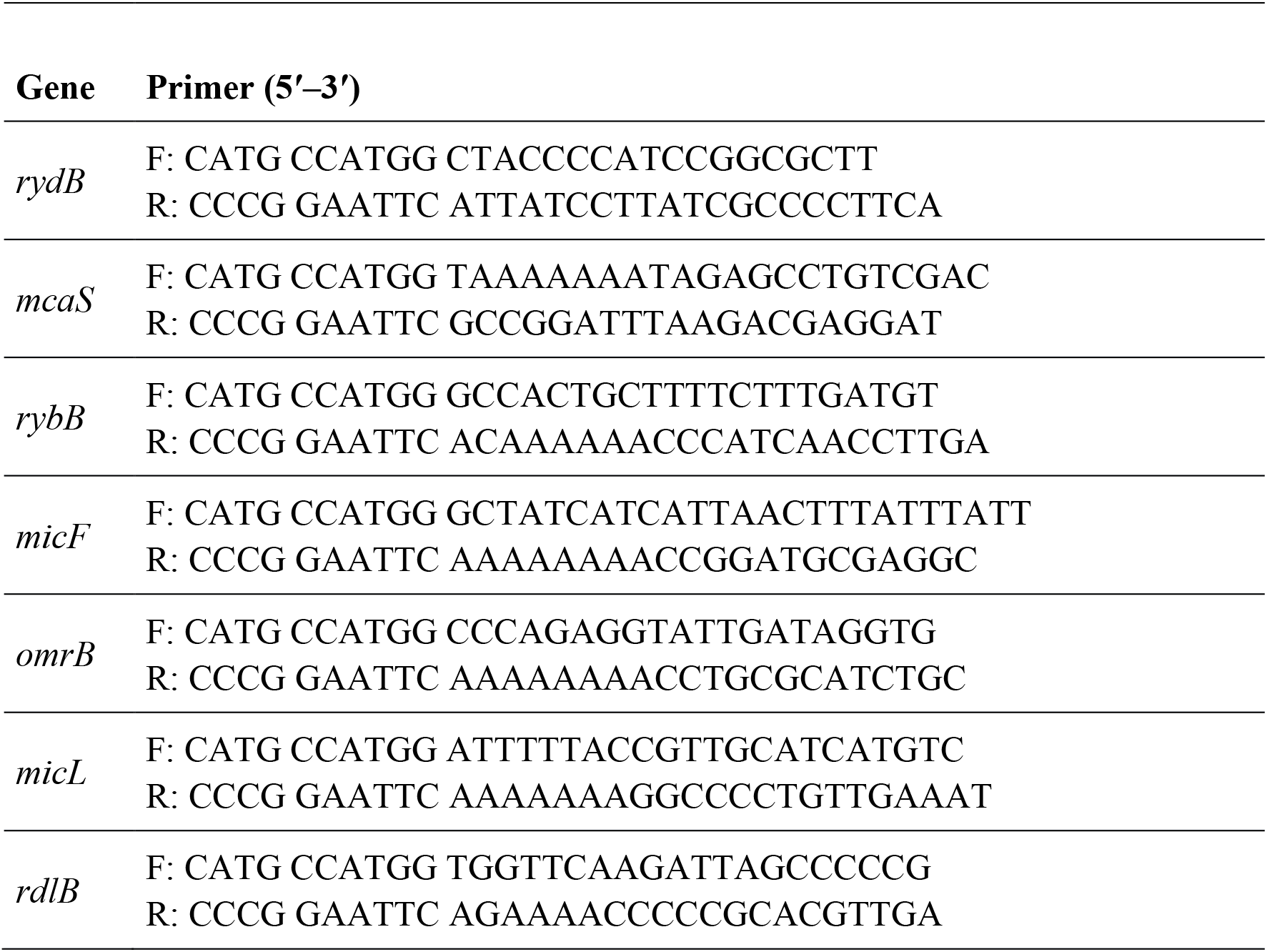
Primers used to generate overexpression strains.

